# A protease-sensing circuit links neutrophil inflammation to virulence regulation in *Streptococcus pyogenes*

**DOI:** 10.64898/2026.05.15.725401

**Authors:** Stephanie Guerra, Ananya Dash, Doris L. LaRock, Christopher N. LaRock

## Abstract

*Streptococcus pyogenes* (Group A *Streptococcus*) causes infections with a disproportionately hyperinflammatory response from the host, such as scarlet fever, necrotizing fasciitis, and toxic shock syndrome. Inflammation is specifically driven by *S. pyogenes* virulence factors, including the protease SpeB, but how inflammation impacts SpeB expression in return during disease is unknown. In this study, we identify a novel interaction between NETosis, a form of inflammatory cell death for neutrophils, and the induction of *speB*. Specifically, while the cathelicidin peptide LL-37 can repress *speB* through the two-component regulatory system CovRS, neutrophil proteases released during NETosis relieve repression of *speB* by degrading another repressor of *speB*, the bacterial protein Vfr. Furthermore, at high cell densities, SpeB autoregulates its expression through similar degradation of Vfr. Abrogating the formation of NETs or depleting neutrophils resulted in *speB* repression *in vivo*, showing the mutual host and pathogen counterattacks collectively lead to the pathological exacerbations characteristic of disease.

## Introduction

*Streptococcus pyogenes* (*Spy*; Group A Streptococcus) is a human-exclusive bacterial pathogen responsible for billions of infections and more than 500,000 deaths annually, making it one of the top ten infectious causes of human mortality worldwide^1^. Most infections are relatively mild, such as impetigo and pharyngitis, but serious, and often fatal diseases like bacteremia, puerperal sepsis, Streptococcal toxic shock syndrome (STSS), and necrotizing fasciitis arise when *Spy* becomes invasive. Virulence factors play a major role in disease progressing by facilitating tissue invasion, evading immune cell killing, and creating intracellular reservoirs^6^.

SpeB is a major virulence factor conserved amongst *Spy*, and is essential for colonization and proliferation within tissue in a variety of infection models^7^. Nonetheless, mutants arise during severe infections that, enigmatically, no longer express *speB*^8,9^. These mutations are most commonly in the two-component system CovRS (CsrRS), which modulates expression of nearly 15% of *Spy* genes, including most virulence factors^10–12^. CovRS senses the host cathelicidin peptide LL-37, produced by epithelial and immune cells as part of innate immune defense and wound repair^5,13–15^. Loss-of-function *covRS* mutations dysregulate their virulence factors in a manner similar to constitutive LL-37 induction, decreasing production of SpeB, while increasing production of capsule (through the *hasABC* operon) and toxins like SLO, NADase, Mac/IdeS, and ^4^. The major inducer of *speB* is RopB in response to the quorum-sensing peptide SIP^16,17^. *In vitro,* the <u>v</u>irulence factor-related (Vfr) protein also appears to antagonize induction, on the basis that *vfr* mutants express greater *speB*^18,19^. How the combination of these possible host and microbial signals are integrated to regulate SpeB during infection is unknown.

This work shows that *Spy* establishes a phenotypically diverse population during infection through the heterogeneous expression of *speB*. Using genetics, RopB, Vfr, and the CovRS system are all found to be essential for generating discrete SpeB-producing and non-producing subpopulations. We show that Vfr is a labile sensor of protease activity that is degraded by SpeB, allowing it to autoregulate it’s own expression by relieving Vfr repression. Furthermore, neutrophil serine proteases also relieve Vfr repression, allowing the bacteria to sense both influx of neutrophils and their activities, with maximal *speB* induction occurring in response to neutrophil extracellular traps (NETs). Despite neutrophils being the major producers of LL-37, which can lead to *speB* repression through CovRS, protease sensing by Vfr ensures SpeB, which is important for resisting neutrophil-produced antimicrobials including LL-37^20^, is still expressed in its presence. Together, our results show how *Spy* navigates the challenge of expressing the right virulence factor at the right time by use of a circuit that integrates host and microbial cues.

## Results

### *S. pyogenes* establishes subpopulation heterogeneity in their expression of *speB*

To examine *speB* expression within *Spy*, we generated a transcriptional fusion of *gfp* with 1000 bp upstream of the *speB* to include P1 and P2 regions of the *speB* promoter^16^. The best-characterized regulator of *speB* is CovRS; *speB* is derepressed when CovR is phosphorylated (CovR∼P), in contrast to the capsule biosynthesis operon *hasABC*, which is induced when CovR is unphosphorylated^12^. CovR phosphorylation depends on the histidine kinase, CovS, which is sensitive to environmental signals. To examine the contribution of *speB* inducers separate from this repressor, we created a tandem reporter that also has the *hasABC* promoter fused to *rfp*^21^ (**Fig. 1A**). To test the accuracy and precision of the reporter construct, *Spy* was grown in concentrations of LL-37 (300 nM) or MgCl_2_ (15 mM) that did not impact bacterial growth (**Fig. S1A**), but which divergently regulate CovR phosphorylation of CovS^12^. As expected for a quorum-sensing regulated protease^17^, GFP (*speB*) is induced as *Spy* approaches late log phase. The addition of LL-37 induced RFP (*hasABC*) and repressed both the expression of GFP (*speB*) (**Fig. 1C**) as well as production of the mature, active protease (**Fig. S1B**). Contrarily, GFP (*speB*) induction was maintained in the presence of MgCl_2_, but RFP (*hasABC*) repressed. Examination by fluorescence microscopy recapitulated these observations, but suggested heterogeneity in these responses within the bacterial population (**Fig. 1B**).

**Fig. 1.**
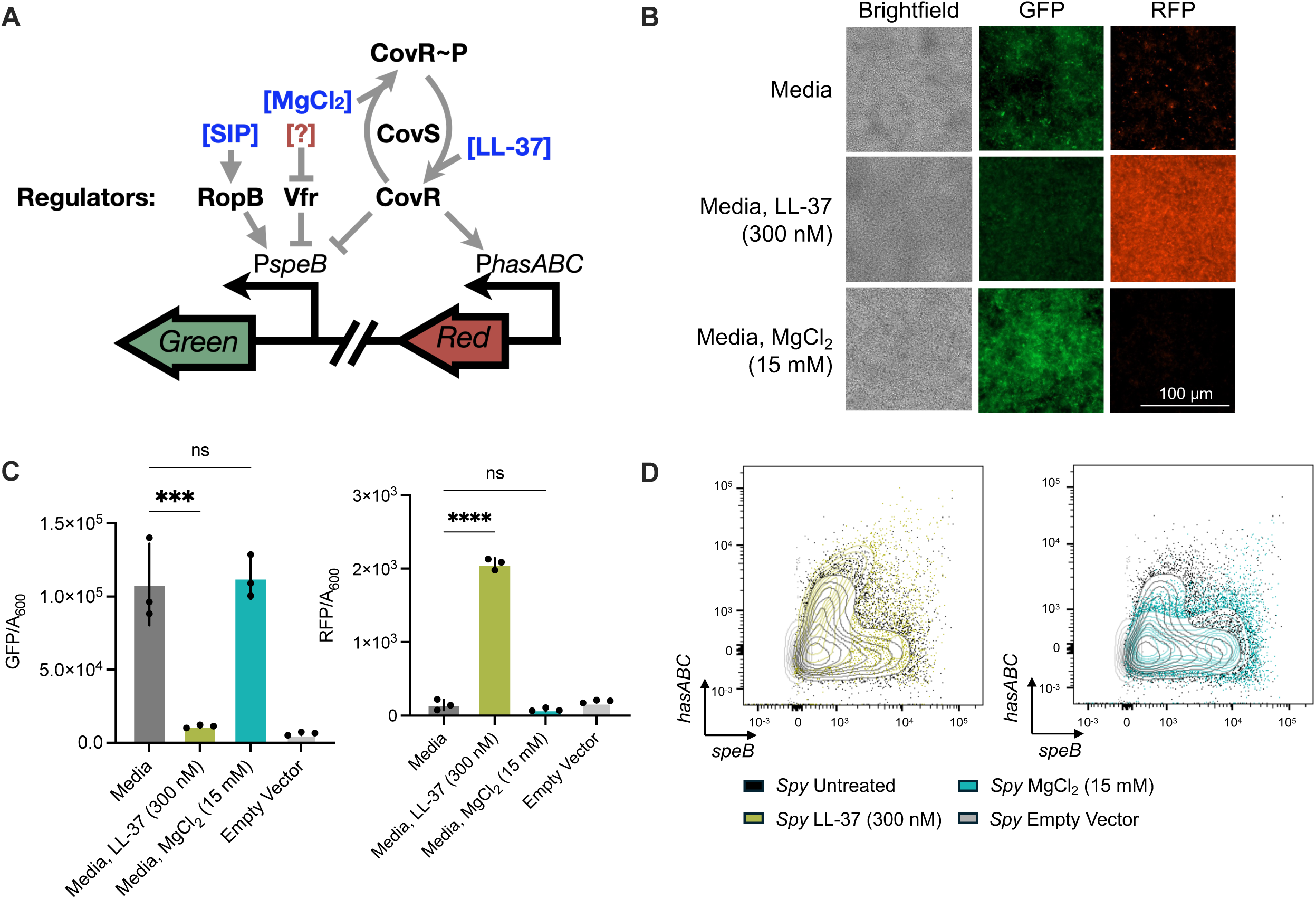
*S. pyogenes* establishes subpopulation heterogeneity in their expression of *speB.* (A) *speB* regulatory circuit schematic. RopB induces *speB* expression in the presence of SIP peptide and Vfr inhibits *speB* expression by blocking RopB-SIP complex. CovS kinase-mediated CovR phosphorylation (CovR∼P) derepresses *speB*, while CovS phosphatase-mediated CovR dephosphorylation represses *speB*. MgCl_2_ and LL-37 mediate CovS kinase and phosphatase, respectively. Induction of *speB* expression was evaluated with GFP fluorescence and *hasABC* expression was evaluated with RFP fluorescence. (B, C, D) Wild-type *Spy* was treated with LL-37 (300 nM) or MgCl_2_ (15 mM). (B) Live-cell fluorescent microscopy with brightfield, GFP, and RFP channels on *Spy* culture grown at stationary phase. Scale bars, 100 µm. (C) Measurement of fluorescence over cell density after 10 h of growth. (D) Flow cytometry demonstrating *speB* expression (GFP; horizontal axis) and *hasABC* expression (RFP; vertical axis) of *Spy* growing at stationary phase. Statistical significance was determined using a one-way ANOVA with Dunnett’s multiple comparisons test. ****P<0.0001, ***P<0.001, ns: not significant.

To quantify this at the single-cell level, we next used flow cytometry. Size, BactoView^TM^ Dead 760/780, and an antibody against the *Spy*-specific antigen Group A Carbohydrate (anti-GAC), were used to gate on single, live cells (**Fig. S1C**). As expected, *Spy* grown with LL-37 had a population shift positive for RFP and negative for GFP (**Fig. 1D**). Conversely, growth with MgCl_2_ resulted in a lower and rightward shift in the population. However, there was significant heterogeneity even in these induced conditions, with large populations of intermediate expression.

Genetic knockouts of known and probable SpeB regulators (**Fig. 1A**) were created to further validate the fluorescent reporters while defining determinants of heterogenicity. None impacted bacterial growth (**Fig. S2A**). A *ΔcovS* mutant constitutively expressed high *hasABC*, (RFP), while repressing *speB*, as expected. A *ΔropB* mutant did not express *speB*, in agreement with its reported importance for *speB* induction^17^. Interestingly, a *ΔspeB* mutant also showed less induction of the *speB* reporter, suggesting the possibility of some autoregulation (**Fig. 2A**). Reporter fluorescence examined by cytometry (**Fig. 2B**) and microscopy (**Fig. S2B**) were consistent with these observations. Similarly, the *ΔspeB* and Δ*covS* mutants expressed slightly less, while *ΔropB* bacteria expressed little (**Fig. 2B, S2B**).

**Fig. 2.**
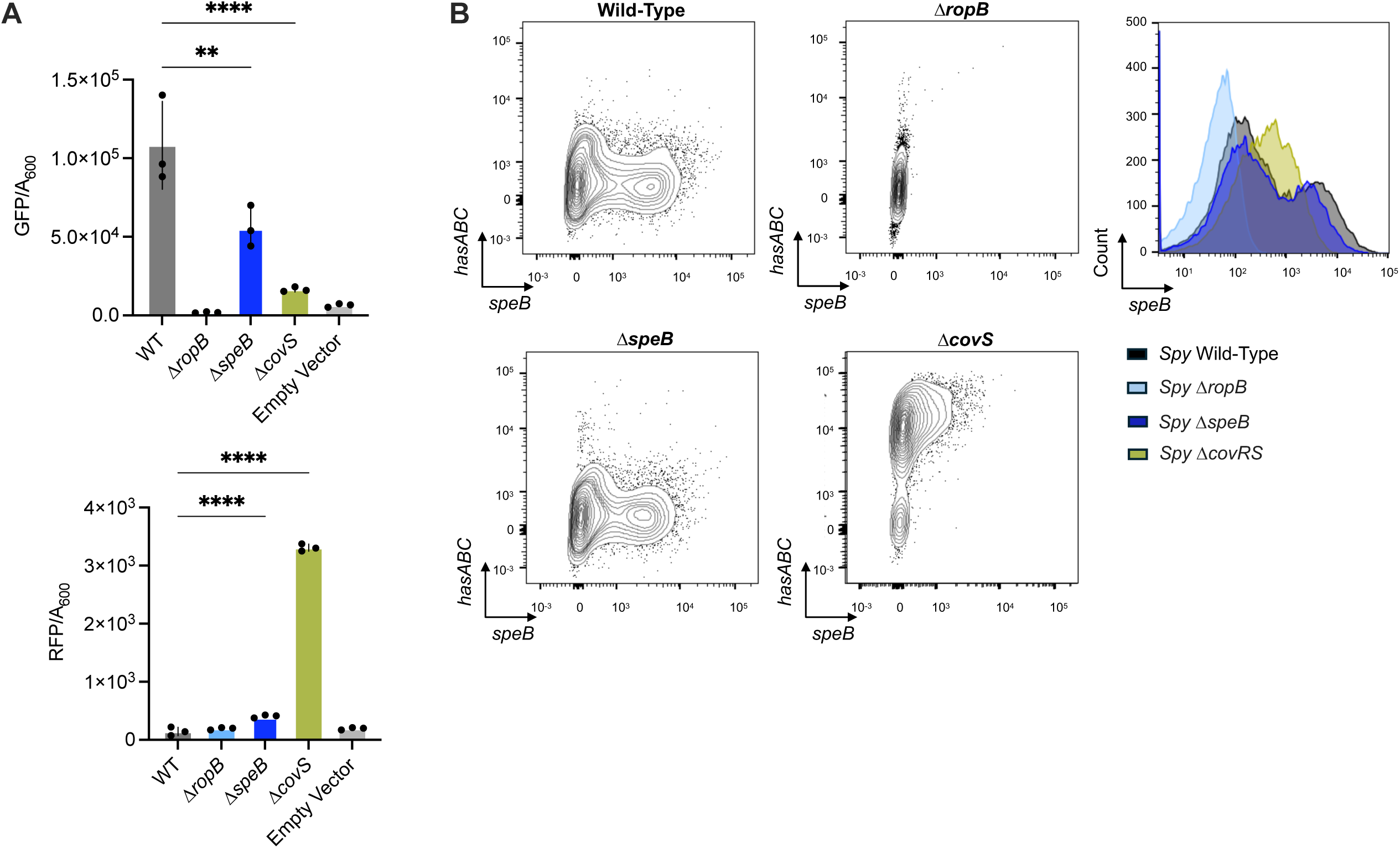
*S. pyogenes* regulators are required for heterogenous *speB* expression. *Spy* genetic control strains of the *speB* regulatory circuit were evaluated. (A) Measurement of fluorescence over cell density after 10 h of growth. (B) Flow cytometry demonstrating *speB* expression (GFP; horizontal axis) and *hasABC* expression (RFP; vertical axis) of *Spy* growing at stationary phase. Statistical significance was determined using a one-way ANOVA with Dunnett’s multiple comparisons test. ****P<0.0001, **P<0.01, ns: not significant.

### Vfr is a dominant repressor for *speB*

Since secreted bacterial factors, including the SIP quorum-sensing peptide, give the potential for bacteria to coordinate expression within the population, we next performed media swap experiments. Wild-type *Spy* was grown to early exponential (EE), mid-late exponential (MLE), or stationary phase (SP) and the spent media removed, filtered, then used to supplement wild-type *Spy* growth carrying the SpeB reporter (**Fig. 3A**). As expected, since the SIP inducer accumulates in stationary phase cultures where SpeB is optimally expressed, SP spent media induced GFP (*speB*) (**Fig. 3B**). Surprisingly, however, growth from earlier growth phase cultures not only lacked *speB* inducer activity but suppressed its expression (**Fig. 3B**). This suggested that these cells produced a soluble factor that dominantly repressed the ability of other cells to produce SpeB. We next repeated the media swap experiment using a *Δvfr* mutant, which highly expressed SpeB (**Fig. S2B**), to see if a soluble factor from wild-type *Spy* could repress this. Spent media from only early-growth phase cultures could significantly repress this (**Fig. 2B**).

**Fig. 3.**
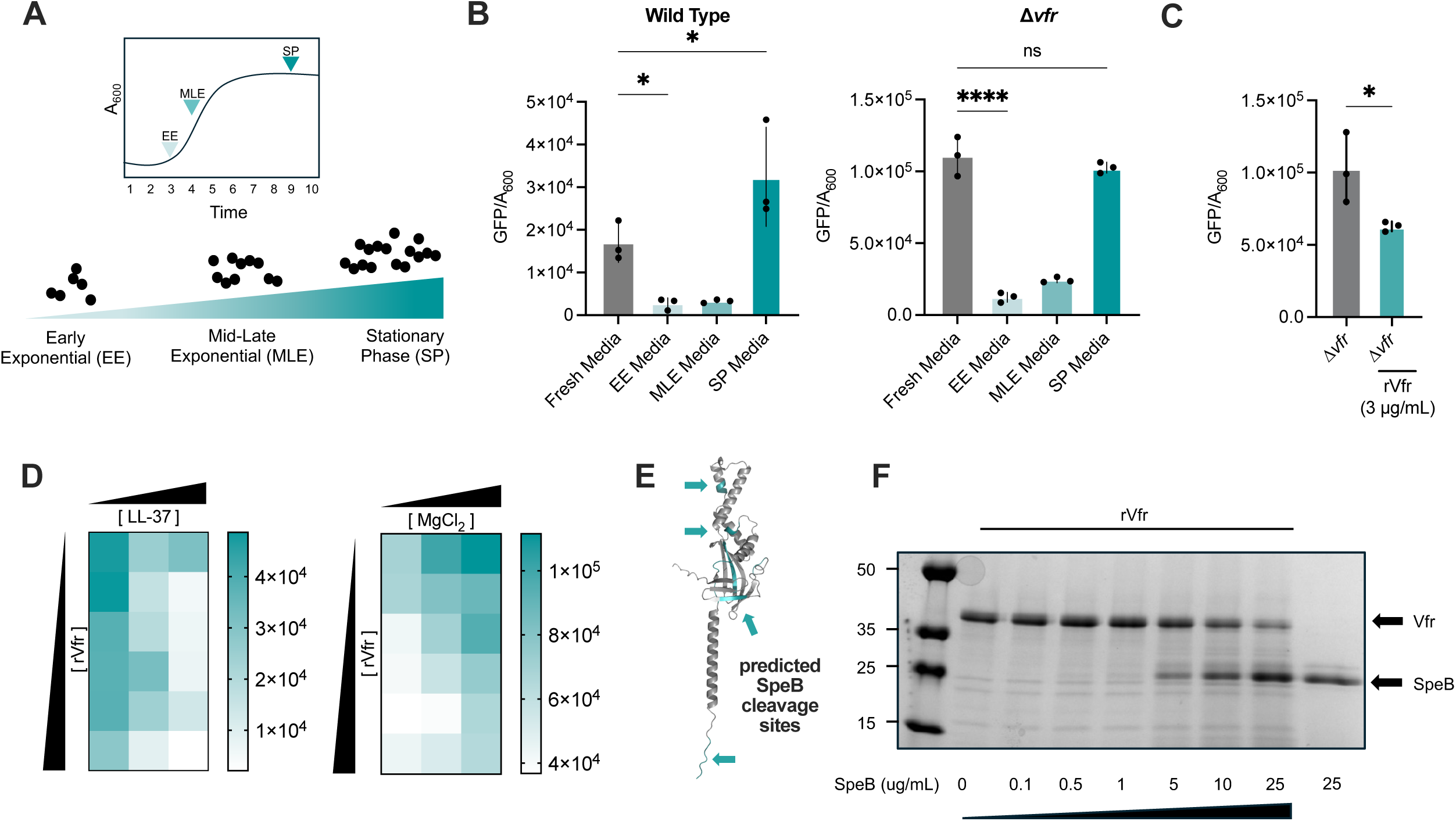
Vfr is a dominant repressor for *speB*. (A, B) Wild-type *Spy* was grown to early exponential (EE), mid-late exponential (MLE), and stationary phase (SP). Spent media from each stage of growth were collected and filter sterilized through 0.2µm filter and inoculated with *Spy* to detect GFP (*speB*) and RFP (*hasABC*) fluorescence. (C) *Spy Δvfr* was grown in the presence or absence of recombinant Vfr (rVfr). (D) Wild-type *Spy* was grown with rVfr (0 - 10 ug/mL) and either LL-37 (0 - 300 nM) or MgCl2 (0 - 15 mM) for 10 h. (E) AlphaFold structure of Vfr with potential SpeB cleavage sites (blue). (F) SDS-PAGE of Vfr (0.3 mg/mL) incubated with incremental concentrations of SpeB for 2.5 h. Statistical significance was determined using a one-way ANOVA with Dunnett’s multiple comparisons test. ****P<0.0001, *P<0.05, ns: not significant.

To assess whether it was Vfr itself in the media suppressing *speB*, we examined the activity of recombinant Vfr (rVfr). rVfr reduced GFP (*speB*) expression significantly in *SpyΔvfr* (**Fig. 3C**). To assess the relative phenotypic dominance of SpeB regulation by autoregulatory mechanisms (Rop-SIP-Vfr dependent) versus environmental signaling mechanisms (CovRS-dependent), wild-type *Spy* containing the dual reporter was grown in the presence of CovRS inducers (LL-37 or MgCl_2_) and Vfr at increasing concentrations. Both LL-37 and Vfr function synergistically in repressing *speB,* while Vfr antagonized MgCl_2_-mediated induction of *speB* (**Fig. 3D**). Notably, *speB* expression in *Spy Δvfr* is unaffected by LL-37 or MgCl_2_, further validating its dominance over CovRS regulation (**Fig. S2C**).

Interestingly, the Vfr structure contains several potential protease SpeB cleavage sites (**Fig. 3E**)^22^. Since degradation could explain the SpeB-dependent changes in GFP (*speB*) expression, rVfr was incubated in the presence and absence of purified SpeB. Vfr degradation was observed in a concentration-dependent manner (**Fig. 3F**). Together, these data highlight that Vfr is a SpeB-labile repressor of SpeB expression.

### *speB* is induced in the presence of immune effectors

Neutrophils are the first line of defense during innate immune response and are quickly recruited during bacterial infections. Neutrophils highly express the immune effector LL-37^5^, which *Spy* recognizes through CovRS, which by current models should repress *speB* expression. *Spy* grown with neutrophil lysates induced *hasABC* expression, consistent with CovRS stimulation, but did not repress *speB* expression (**Fig. 4A**). Similarly, in the mouse invasive infection model and in whole human blood, *Spy* induced *hasABC* while populations of high *speB* expressors were maintained (**Fig. 4B**). *Spy* with regulator knockouts were also evaluated to determine consistency in mouse and blood tissue infections with *in vitro* regulation. *Δvfr* maintained high *speB* expression, *ΔropB* bacteria were negative for *speB* expression, and *ΔcovS* contained a *hasABC* high population (**Fig. S3A**).

**Fig. 4.**
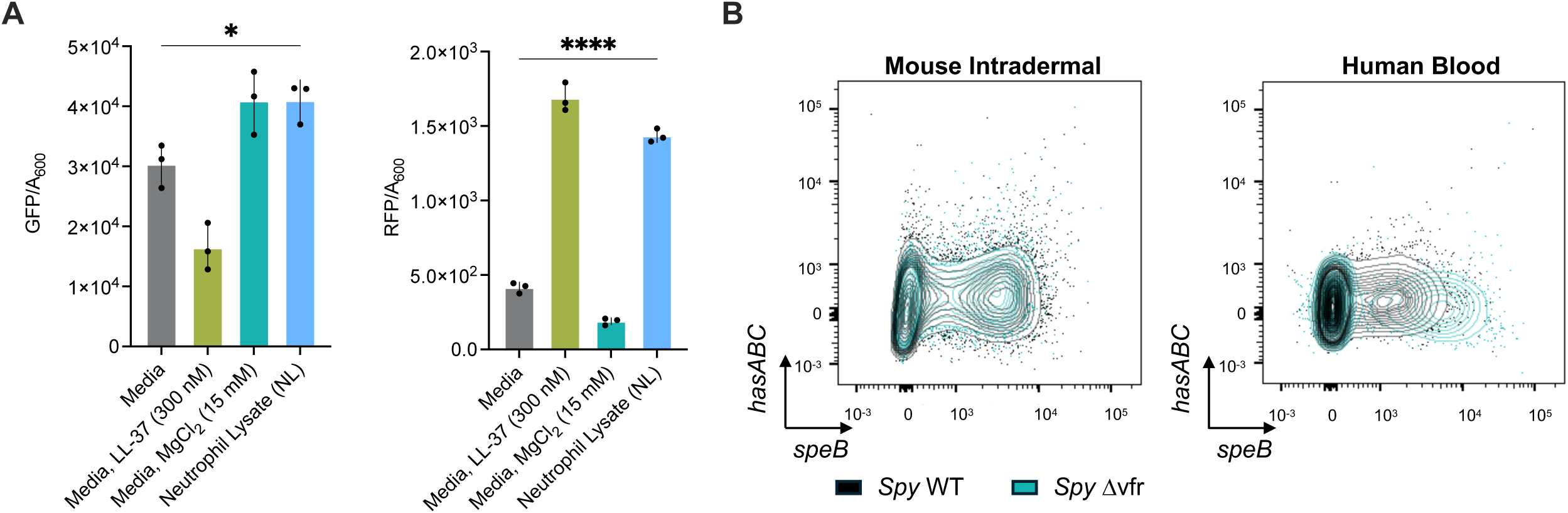
*speB* expression is induced in the presence of immune effectors. (A) Evaluating the effects of neutrophil lysate on *speB* and *hasABC* expression. *Spy* grew in the presence of neutrophil lysates (10^6^ cells/mL) compared to *Spy* growing in RPMI 5% THY. Treatment with LL-37 (300 nM) and MgCl_2_ (15 mM) served as controls. (B) Flow cytometry demonstrating *speB* (GFP fluorescence; horizontal axis) and *hasABC* induction (RFP fluorescence; vertical axis) of 10^8^ CFU of *Spy* during mouse intradermal and human blood infections after 24 h and 4 h, respectively.

### Immune effectors induce *speB* through Vfr

This high-level of expression *in vivo* suggested that *speB* could be induced, not just repressed, in the presence of immune effectors. Along with LL-37, neutrophil granules store many major inflammatory components secreted during infections^23,24^. To assess whether other neutrophil derived molecules play a role in *speB* regulation, neutrophil lysates were fractionated and examined for their ability to induce or repress *speB* expression. Upon supplementation, GFP (*speB)* was induced by three fractions (10-12) containing active proteases (**Fig. 5AB**).

**Fig. 5.**
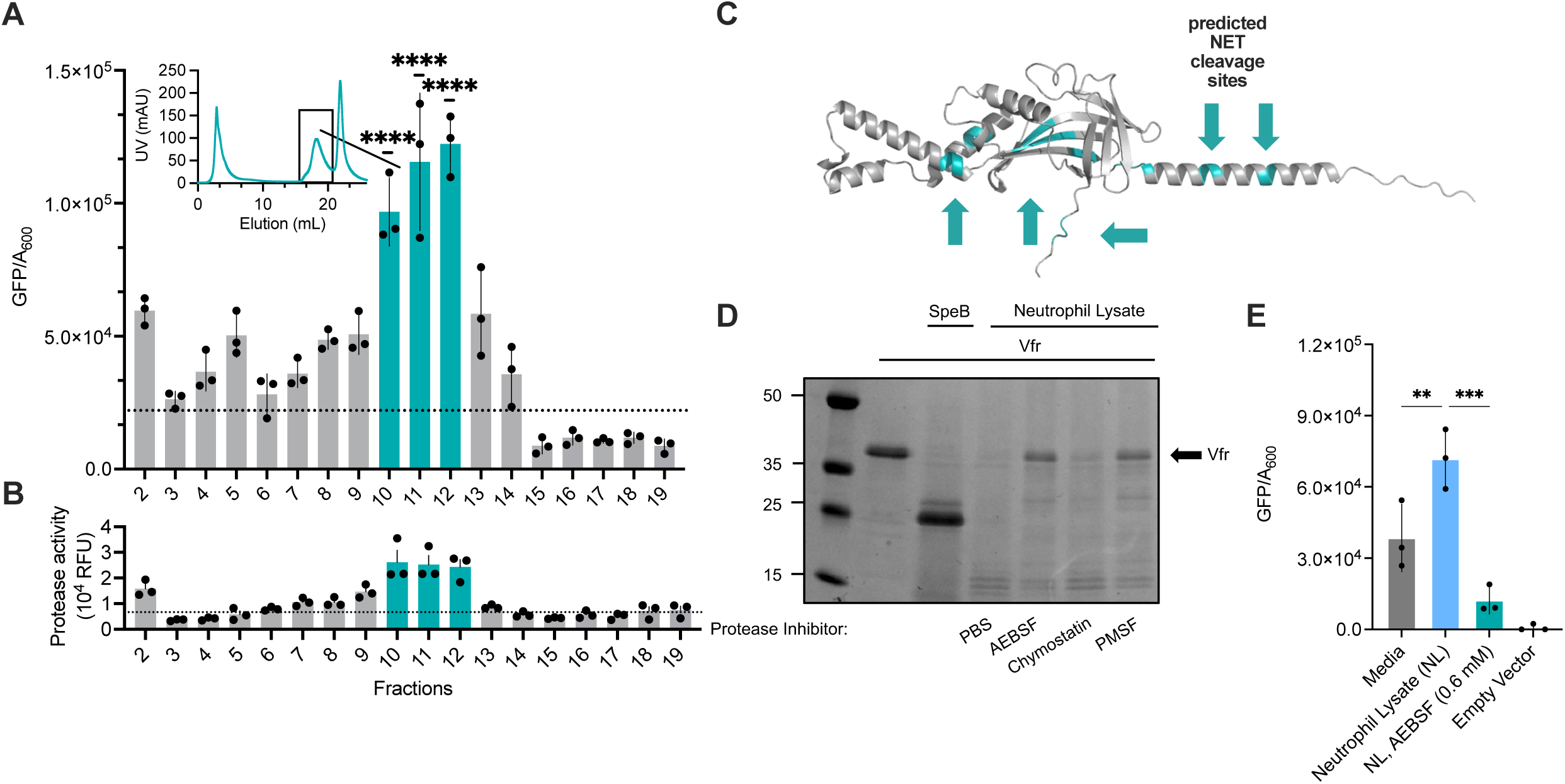
Immune effectors induce *speB* through Vfr. (A, B) Neutrophil lysates were fractioned based on net surface charge through anionic exchange. (A) Fractions were used to supplement *Spy* growth and *speB* expression was evaluated with GFP fluorescence at late log phase. Protein content within each elution (mL) was detected by UV (mAU) (Upper left). (B) Fractions were also used to evaluate protease activity. (C) AlphaFold structure of Vfr with potential NET protease cleavage sites (blue). (D) SDS-PAGE of Vfr (0.3 mg/mL) incubated with lysate from neutrophils (10^6^ cells/mL) treated with inhibitors AEBSF, Chymostatin, or PMSF. PBS served as a vehicle control. (E) *Spy* grew in the presence of neutrophil lysates (10^6^ cells/mL) and AEBSF (0.6 mM) compared to *Spy* growing in RPMI 5% THY. Statistical significance was determined using a one-way ANOVA with Dunnett’s multiple comparisons test. ****P<0.0001, ***P<0.001, **P<0.01.

While *Spy* is resistant NET killing^25,26^, the immune effectors released by neutrophils during NETosis, include not only LL-37, but proteases like neutrophil elastase (NE)^27^. Based on the protease profile of NETs^28^, Vfr contains several potential sites for cleavage (**Fig. 5C**). Neutrophils degraded rVfr (**Fig. 5D**), as did purified NE (**Fig. S3C**). The NE-specific inhibitor, BAY-678, was not sufficient to inhibit neutrophil lysate-mediated rVfr degradation, but serine inhibitors AEBSF and PMSF did (**Fig. 5D, S3C**), suggesting a sufficiency but not requirement for other neutrophil proteases. Moreover, *Spy* grown with neutrophil lysates and inhibitor AEBSF had resulted in *speB* inhibition expression, compared to neutrophil lysate alone (**Fig. 5E)**. Notably, these conditions maintained *hasABC* expression, consistent with CovRS stimulation (**Fig. S3D)**. Altogether, these data suggest Vfr is a substrate for both bacterial and host-mediated degradation.

### SpeB and neutrophils are sufficient to induce *speB* expression

To examine the impact that neutrophils have on *speB* regulation *in vivo,* neutrophils were depleted by anti-Ly6G treatment, as previously^29^, significantly decreasing their numbers (**Fig. S4A**). After a 24 h intradermal infection, *Spy* from the tissue was examined by flow cytometry for *speB* expression, relative to a *Δvfr* control to define the high-expressing population (**Fig. S4B**). *Spy* expression of *speB* was only moderately decreased in the absence of neutrophils (**Fig. 6B**). However, infection of neutropenic mice with a *ΔspeB*, where there is neither host or microbial proteases to cleave Vfr, prevented *speB* expression (**Fig. 6B**). Mice deficient in peptidyl-arginine deiminase 4 (PAD4) are unable to form NETs^30^. To separately examine NET contributions *PAD4^-/-^* mice were infected with *Spy* and *speB* expression examined. SpeB expression was reduced in *PAD4^-/-^* mice, additively with a contribution of SpeB itself (**Fig. 6C**). Collectively, this data suggests that *Spy* integrates the sensing of both neutrophils and feedback from SpeB to titrate expression during infection.

**Fig. 6.**
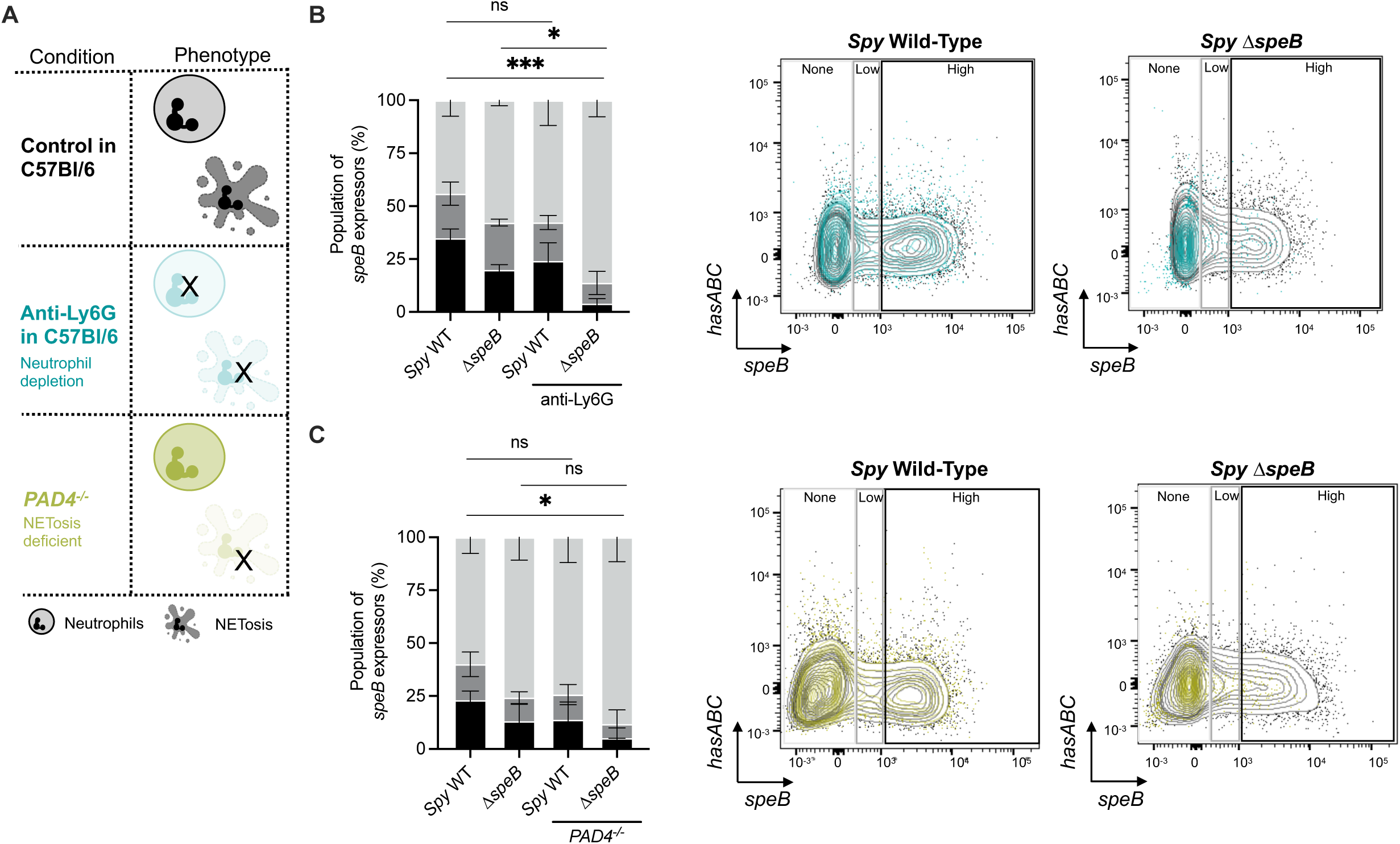
SpeB and neutrophils are sufficient to induce *speB* expression. (A) Diagram of mouse intradermal infection model in three different conditions: neutrophil-depleted mouse (anti-Ly6G treatment); NET-deficient mouse (*PAD4^-/-^*); and neutrophil competent (wild-type) control. (B, C) Flow cytometry on mouse skin lesions from 24 h intradermal infection of 10^8^ CFU of *Spy* or *Spy ΔspeB*. Population of *speB* expressors was determined based on fluorescent intensity during flow cytometry. No expressors range was based on the negative empty vector control. (B) C57Bl/6 mice were treated with neutrophil-depleting antibody, anti-Ly6G, (blue) or control (black) for 24 h then infected. (C) NETosis-deficient mice (*PAD4^-/-^*, yellow) or control (C57Bl/6 mice, black) were infected. Statistical significance of high *speB* expressors was determined using a two-way ANOVA with Dunnett’s multiple comparisons test. ***P<0.001, *P<0.05, ns: not significant.

## Discussion

Since a bacterium must adapt to environmental changes, they modulate expression in response to diverse stimuli, including stress, metabolites, quorum sensing, pH, and temperature. The niche of *Spy* is exclusively the human, potentially limiting its exposure to the range that some other species face. Yet within this host, it is nonetheless presented with challenges. Foremost amongst these are the immune defenses, which escalate through the course of infection. Fine-tuning its regulation toward the virulence factors needed to overcome this specific challenge is key to the species. SpeB is a potent pro-inflammatory virulence factor required to establish infection and has key roles in determining the course and severity of infections^6^. Accordingly, the regulation of SpeB must be tied to circumstances where it will be advantageous. Regulation of *speB* is thus multifactorial, as it integrates sensors to promote colonization and survival. In this study, we highlight a novel mechanism for SpeB regulation through Vfr detection of neutrophils and their activation state. Our data shows that neutrophils, despite functioning as an LL-37 reservoir expected to repress *speB* expression through the CovRS system, actually induces it during invasive skin infections.

The relative contributions of these regulators could vary by disease^31^. In response to inflammation, neutrophils are recruited to infection sites to engulf and inactivate pathogens. Despite neutrophils playing a major role in combating most infections, several studies indicate they can worsen *Spy* infections, in particular, in the upper respiratory tract^29,32^. Here, RopB can detect the SIP homologs used for quorum sensing by other *Streptococci* species of the upper respiratory microbiota^33^. This ensures SpeB expression at low cell densities and could allow SpeB expression even before neutrophil infiltration. Importantly, the type of CovS mutants that interfere with SpeB expression do not occur during human pharyngitis or experimental mouse models of it, and *ΔspeB* mutants are highly attenuated, indicating its essentiality at this site^29^. This would subsequently require SpeB expression during infections, suggesting repression by LL-37, released by the abundant neutrophils recruited during infection, is, at best, limited.

Our work suggests that Vfr functions as a biosensor of protease activity that regulates the major protease of *Spy*, SpeB. It is additionally degraded by neutrophil proteases, thereby relieving repression in the presence of these important immune cells. Originally identified through a transposon mutagenesis screen^18^, Vfr has been suspected to regulate SpeB in a RopB-dependent manner^19^. Our results showing that Vfr functions as a negative regulator of *speB* early during growth in culture is consistent with this, but the mechanism in which SIP quorum sensing relieves Vfr-mediated repression at late and stationary growth remains to be elucidated. Since the SpeB protease itself can degrade Vfr, autoregulation may play a role in this temporal regulation. However, it is not clear whether this would be relevant during infection, since the host can provide the proteases to overcome this. Similarly, CovRS mediates repression of SpeB in some *in vitro* conditions, such as with LL-37 alone. However, since repression is not maintained in the presence of the major LL-37-producing cells, neutrophils, the effect may be more limited during infections, either due to heterogeneity in the population or dominant effects of Vfr. Importantly, other than releasing LL-37, neutrophils undergoing NETosis or degranulating release proteases, including NE^34^, that degrade Vfr. Together, this highlights the importance of *in vivo* data for examining pathways such as the regulation of SpeB, since pathways sensing multiple bacterial and host factors can intersect.

In summary, we demonstrate how *Spy* overcomes challenges in the temporal regulation of the virulence factor SpeB by a circuit that integrates multiple considerations in the host-pathogen interactions. Clear Vfr homologues can be found throughout the Streptococci (*S. agalactiae, S. urinalis, S. pseudoporcinus, S. porcinus, S. uberis, S. iniae, S. equi, S. dysgalactiae, S. canis*), despite none of these species encoding a SpeB homolog. This suggests that Vfr may be a regulator module more generally, and that through these other species senses broader factors as well. We propose a model wherein *Spy*, through the Vfr protein, coordinates expression of SpeB within the bacterial population. At high cell densities, quorum sensing is sufficient to induce expression. However, Vfr allows expression at lower cell densities if there is sufficient protease to degrade it. This can be accomplished if there are some SpeB-expressing cells already, thus leading to further coordination between cells independent of classic quorum-sensing, or, an override in the instance of proximate protease-producing immune cells. This further allows heterogeneity within the population, as each cell is exposed to different concentrations of SIP, LL-37, VFR, and protease. Altogether, beyond demonstrating a previously undescribed mechanism for regulating a protease through a natural biosensor for its activity, we show that this integrated regulation poises the pathogen to respond to shifts in innate immune pressure as part of the sophisticated virulence strategy of *Spy*.

## Resource Availability

### Lead contact

Requests for further information and resources should be directed to and will be fulfilled by the lead contact, Christopher LaRock

### Materials availability

All unique reagents generated in this study are available from the lead contact upon completion of a material transfer agreement.

### Data and code availability

This study did not generate any original sequence data or code. Any additional information required to reanalyze the data reported in this study is available from the lead contact upon request.

## Acknowledgements

We thank Victor Nizet and Andrew Varble for strains, the anonymous blood donors, and members of LaRock lab for helpful discussions. This work was supported in part by the Emory University Integrated Cellular Imaging Core Facility (RRID:SCR_023534), the Emory Flow Cytometry Core (EFCC) Facility (RRID:SCR_023536), and the Children’s Healthcare of Atlanta and Emory University’s Children’s Clinical and Translational Discovery Core. Additional support was provided by the National Center for Georgia Clinical & Translational Science Alliance of the National Institutes of Health under Award Number UL1TR002378. This work was supported by the National Institute of Allergy and Infectious Diseases of the NIH under award numbers AI153071 (C.N.L.) and AI180089 (C.N.L.), training grants AI106699 (S.G.), AI179103 (S.G.), an American Heart Association Predoctoral Fellowship 25PRE1372818 (A.D.), and a Burroughs Wellcome Fund Investigator in the Pathogenesis of Infectious Disease award (C.N.L.). The content is solely the responsibility of the authors and does not necessarily reflect the official views of the National Institutes of Health.

## Author Contributions

Conceptualization, C.N.L; investigation and analysis, S.G., A.D., D.L.L, C.N.L.; writing of the original draft, S.G. and C.N.L.; reviewing and editing, S.G. and C.N.L.; funding, S.G., A.D., C.N.L.; supervision, C.N.L.

## Supporting Information Legends

**Fig. S1.** (A) Wild-type *Spy* growth kinetics determined through optical density at 600 nm (O.D. 600) in RPMI, 5% THY with LL-37 (300 nM) or MgCl_2_ (15 mM). (B) SpeB activity was measured using the fluorescent peptide sub103. (C) Flow cytometry gating strategy for measuring *Spy* GFP and RFP fluorescence. Samples were selected based on particle size (FSC-Area, SSC-Area), then selected for into single cells (FSC-Area, Height; SSC-Area, Height). Sample containing Group A Carbohydrate (APC-A positive population; right peak) were selected. Lastly, live cell population was selected (APC-Cy7-A negative population; left peak). (D) Flow cytometry demonstrating *speB* expression (GFP; horizontal axis) and *hasABC* expression (RFP; vertical axis) of *Spy* growing at stationary phase separated in panels based on treatment.

**Fig. S2.** (A) *Spy* growth kinetics determined through optical density at 600 nm (O.D. 600) in RPMI, 5% THY. (B) Lionheart live-cell fluorescent microscopy with brightfield, GFP, and RFP channels on of *Spy* culture grown at stationary phase. Scale bars, 100 µm. (C) Measurement of fluorescence over cell density after 10 h of growth. Wild-type and Δ*vfr Spy* were treated with LL-37 (300 nM) or MgCl_2_ (15 mM).

**Fig. S3.** (A) Regulation of *speB* and *hasABC* during mouse intradermal and human blood infections. Flow cytometry demonstrating *speB* (GFP fluorescence; horizontal axis) and *hasABC* induction (RFP fluorescence; vertical axis) of 10^8^ CFU of *Spy strains.* (B) Colony Forming Units (CFU) of *Spy* within 4 h human blood infection was measured by plating. (C) SDS-PAGE of Vfr (0.3 mg/mL) incubated with recombinant Neutrophil Elastase (rNE) or neutrophil lysate (10^6^ cells/mL) with inhibitor BAY-678. (D) Measurement of RFP fluorescence (*hasABC*) over cell density after 10 h of growth. (E) *Spy* growth kinetics determined through optical density at 600 nm (O.D. 600).

**Fig. S4.** (A) Flow cytometry gating strategy for neutrophil depletion model with anti-Ly6G. Singlets were selected through side (SSC-H, SSC-W) and forward (FSC-H, FSC-W) scatter. Population of live cells were selected based on BUV480-A fluorescence (left). Granulocytes were selected based on the presence of CD45, and neutrophils were identified based on presence of Ly6G and CD11. (B, C) Flow cytometry demonstrating *speB* (GFP fluorescence; horizontal axis) and *hasABC* induction (RFP fluorescence; vertical axis) of 10^8^ CFU of *Spy. Spy*Δ*vfr* strain in anti-Ly6G neutrophil depletion model (B). Control strains, *Spy* empty vector and *Δvfr*, during mouse intradermal infection with *PAD4*^-/-^(C).

## Methods

### Bacterial strains and growth conditions

All strains are described in **Table 1**. *Spy* strains were routinely grown in Todd Hewitt broth with 5% yeast (THY) at 37°C with 5% CO_2_. Bacterial aliquots were washed in PBS and resuspended in PBS with 20% glycerol for storage at −80°C and grown fresh for each experiment. The bacterial mutations *ΔcovS, ΔropB, and Δvfr* were obtained through lambda red recombineering, as described previously^39^. Briefly, a kanamycin resistance cassette was PCR amplified with the primer sequences outlined in **Table 2**, each carrying 5’ homology to the chromosomal sequence flanking the sites of the desired mutation. *Spy* 5448 carrying recombineering plasmid pAV488 were electroporated with the PCR product and selected for kanamycin resistance. Curing of the recombineering plasmid was achieved by selecting colonies susceptible to chloramphenicol. Gene knockout and lack of spurious secondary-site mutations were validated by whole-genome sequencing (Plasmidsaurus). Reference sequences and plasmids are detailed in **Table 3**.

**Table 1:** Strain List.

| *S. pyogenes* strain | Description | Reference |
| --- | --- | --- |
| M1 5448 | Wild-type M1 | Ref<sup>35</sup> |
| 5448ΔcovR | Deletion of *covR* | This study |
| 5448ΔcovS | Deletion of *covS* | This study |
| 5448ΔropB | Deletion of *ropB* | This study |
| 5448ΔspeB | Deletion of *speB* | Ref<sup>36</sup> |
| 5448Δvfr | Deletion of *vfr* | This study |

**Table 2:**
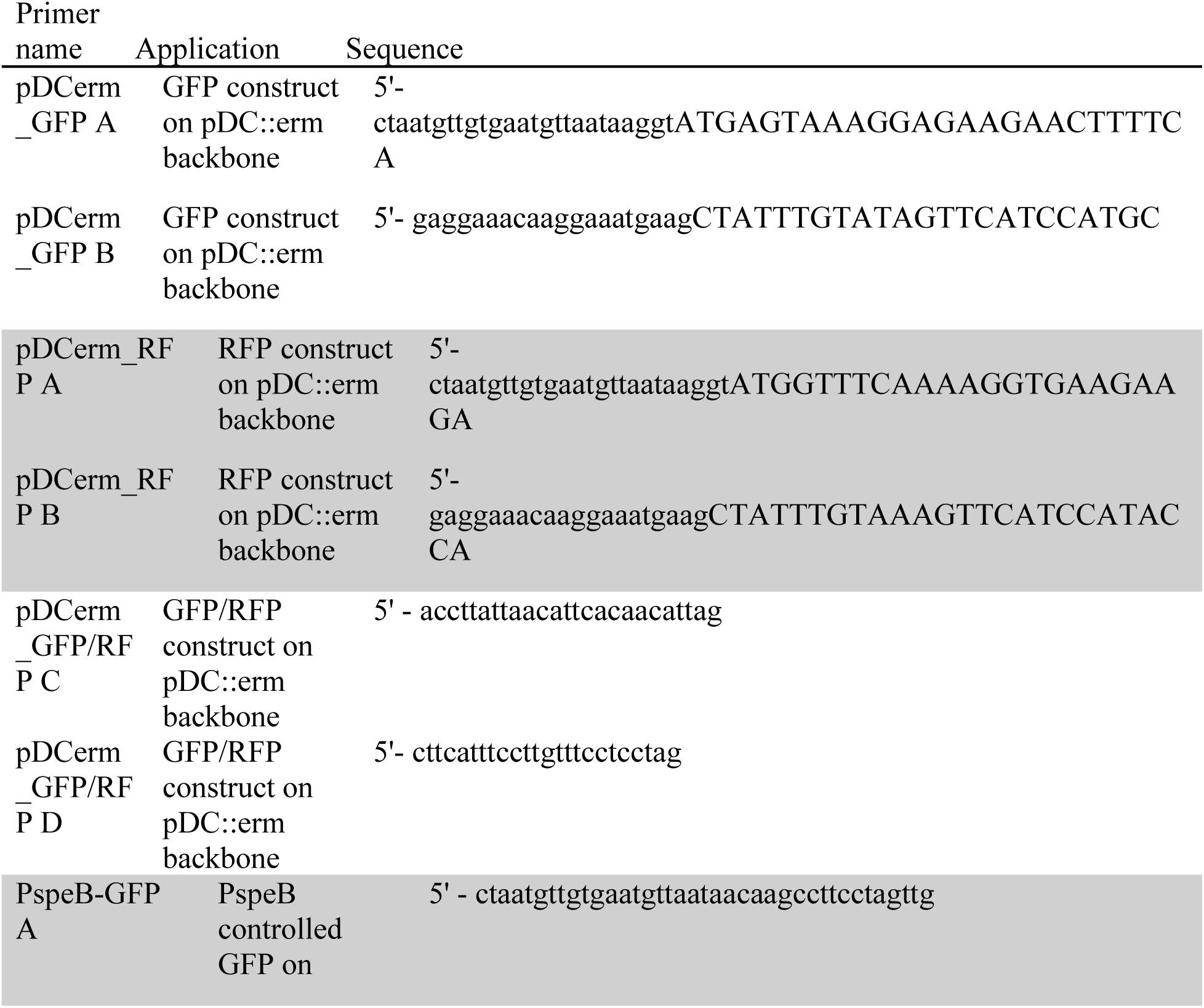
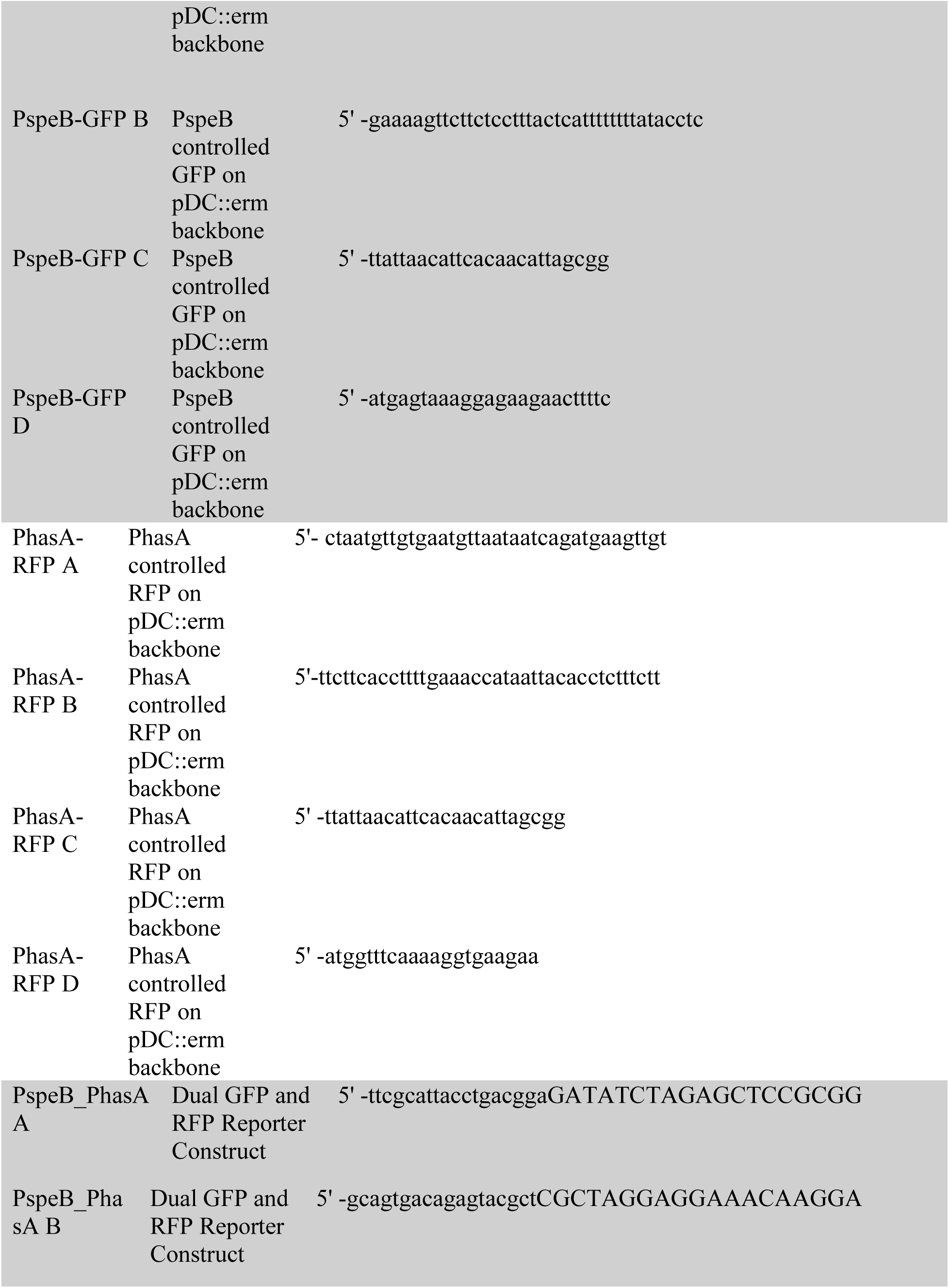
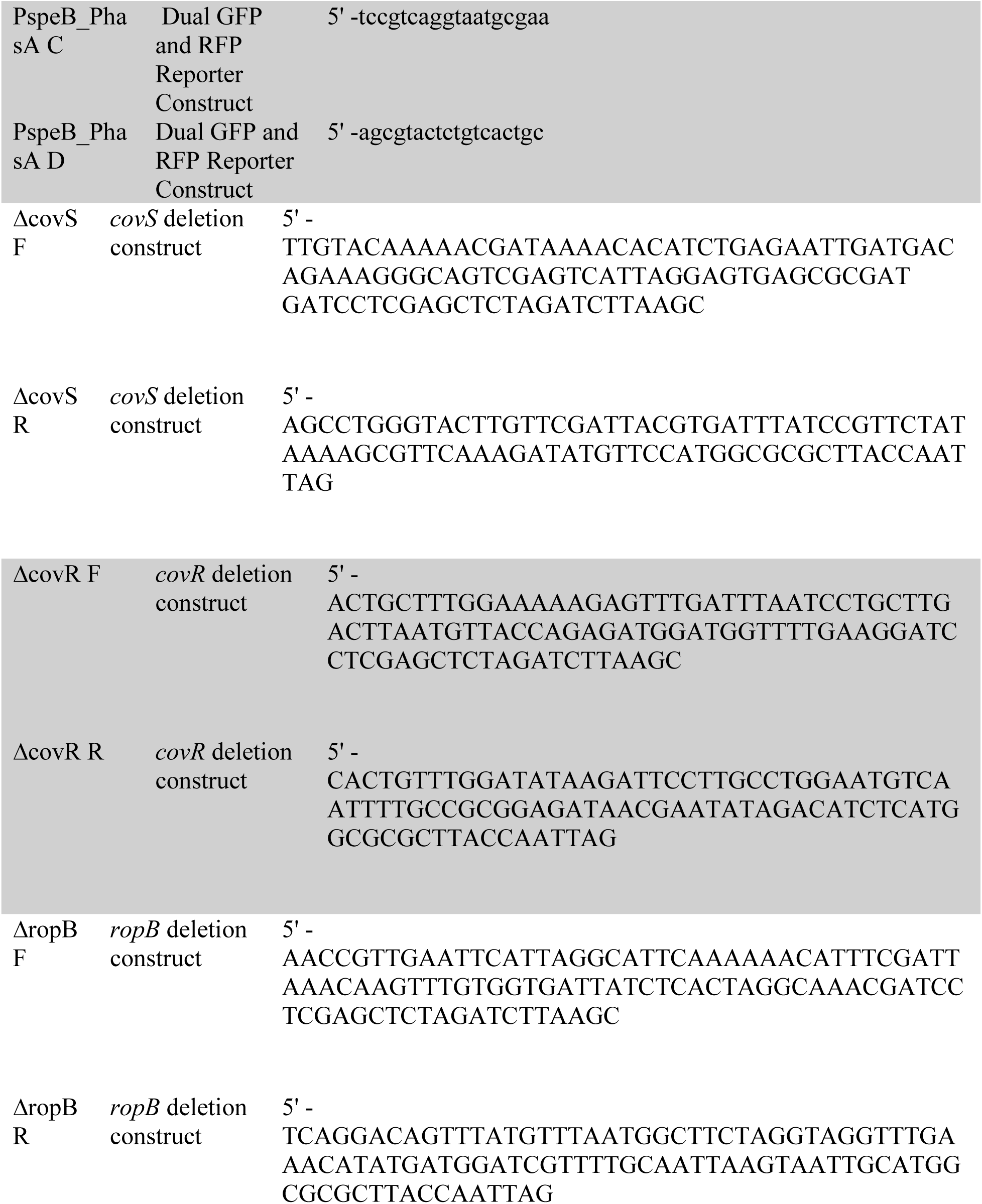
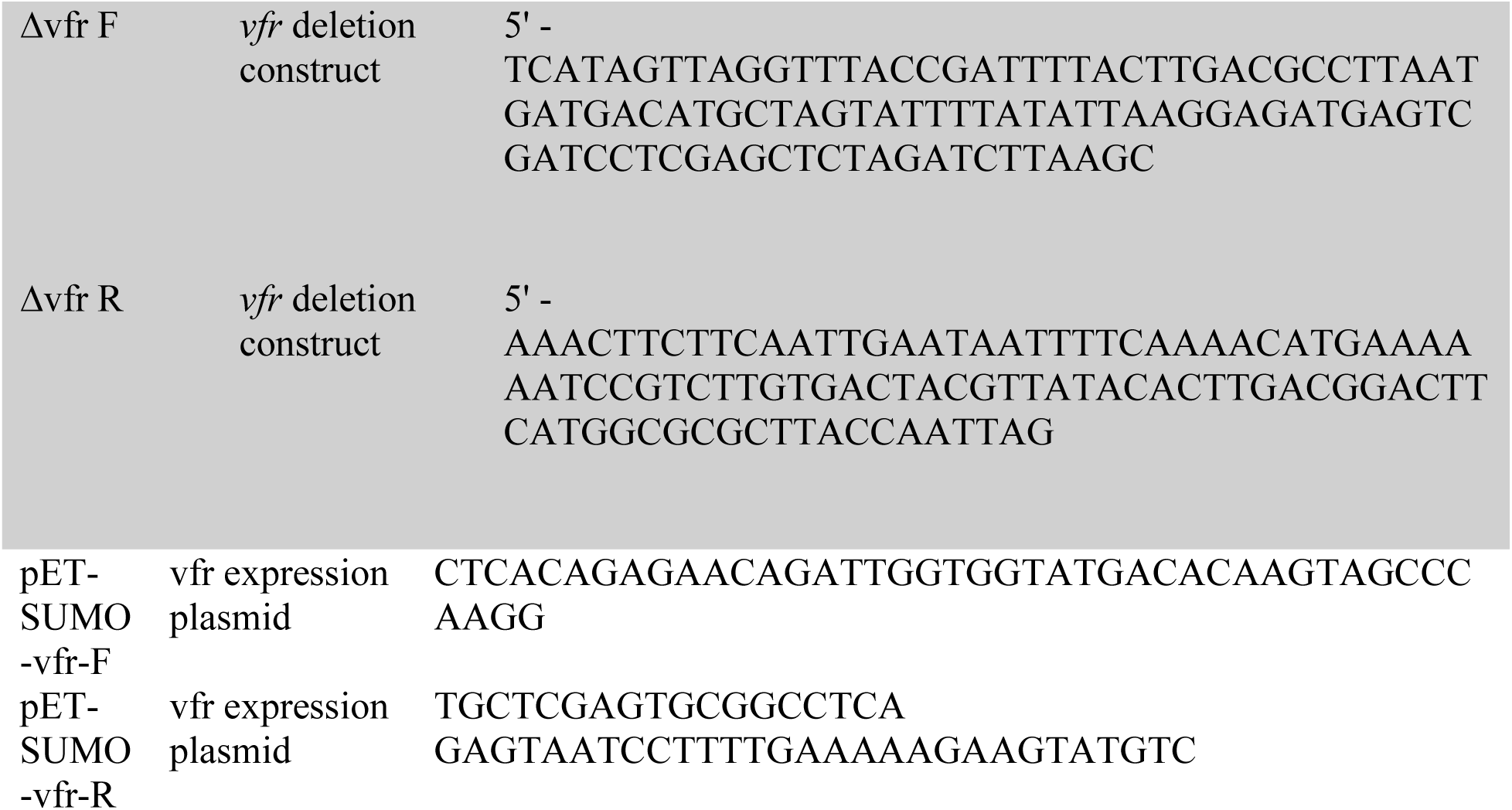
Cloning Primers Primer.

**Table 3:** Plasmids.

| Plasmid name | Description | Reference |
| --- | --- | --- |
| pDCerm | Reporter backbone | Ref<sup>37</sup> |
| pGFP | Plasmid with *gfp* template | GenBank: OM212390.1 |
| pRFP | Plasmid with *rfp* template | GenBank: KM521211.1 |
| pDCerm GFP | Reporter construct | This study |
| pDCerm mCherry | Reporter construct | This study |
| pDCerm-PspeB::GFP | *speB* expression reporter | This study |
| pDCerm-PhasA-mCherry | *hasA* expression reporter | This study |
| pDCerm-PspeB::GFP-PhasA::mCherry | Dual *speB* and *hasA* expression reporter | This study |
| pET-SUMO | Expression vector | Ref<sup>38</sup> |
| pET-SUMO Vfr | Vfr expression vector | This study |
| pAV488 | Recombineering plasmid | Ref<sup>39</sup> |

### Plasmids

Fluorescent reporter plasmids pDCerm-*PspeB::GFP,* pDCerm*-PhasA::RFP*, and pDCerm-*PspeB::GFP-PhasA::RFP* were created for this study by Polymerase Incomplete Primer Extension (PIPE) cloning technique, as previously described^40^. *Spy* promoters were amplified from 5448, GFP from (GenBank: OM212390.1), and RFP from (GenBank: KM521211.1) for insertion into pDCerm^41^ using the primers in **Table 2**. The sequence was validated using whole plasmid sequencing (Plasmidsaurus) and constructs transformed into each *Spy* strain by electroporation. Expression vector pETxSUMO-Vfr was created using primers Vfr F and Vfr R to amplify *vfr* from GAS 5448 and the previously described PIPE primers pETxSUMO F and pETxSUMO R to amplify the expression vector^40^. Reference sequences and plasmids are detailed in **Table 3**.

### Fluorescence during growth

*Spy* 5448 strains grown in Todd-Hewitt Broth with 5% yeast (THY) to mid-exponential phase were used to inoculate fresh, phenol red-free RPMI supplemented with THY (5%) and erythromycin (2 µg/mL) in a 96-well black, clear-bottom plate (Costar). Cultures were grown for 10 hours at 37 °C and 5% CO_2_ with measurements of absorbance (600 nm), GFP (ex. 479 nm, em. 520 nm), and RFP (ex. 579 nm, em. 616 nm) using a BioTek Synergy H1 plate reader. Expression of *speB* and *hasABC* were analyzed by fluorescence of GFP or RFP over absorbance. Supplementation with LL-37 300 nM (GeneScript) and MgCl_2_ 15 mM (Sigma M9272).

### Whole blood and neutrophil experiments

Whole blood was collected from healthy adult donors with informed consent and approval from the Emory University’s Children’s Clinical and Translational Discovery Core. For infections, 10^8^ CFU bacteria suspended in 100 μL of PBS and used to inoculate 400 µl of whole blood. Inoculated blood was incubated on a rotisserie mixer for 4 h. After 4 h, to lyse host cells, the inoculum was treated with Triton X 0.05% (Sigma T8787) for 15 minutes, then samples were stained and fixed for analysis. For neutrophil experiments, neutrophils were isolated from whole human blood by centrifugation in PolymorphPrep (AxisShield) as previously^42^, then diluted in RPMI. To obtain protein content from neutrophils, suspension with neutrophils in RPMI were lysed via sonication (11% amplitude for 4 minutes at 30 second intervals) and centrifuged at 6,000 x g for 5 minutes to remove cellular debris. Neutrophil lysates were used for plate reader analysis or SDS PAGE. Serine protease inhibitor AEBSF 0.6 mM (Sigma 508436) was used for plate reader analysis.

### Animal experiments

All animal use and procedures were performed with approval from Emory Institutional Animal Care and Use Committee. Mice were housed in specific pathogen-free conditions with a 14 h light/10 h dark cycle in a standard ambient environment (∼20 °C and ∼50% humidity) in ABSL-2 conditions. Experiments were performed using both male and female wild-type C57Bl/6 and NET-deficient *PAD4^-/-^*(JAX #030315)^30^ mice of 8-12 weeks of age (Jackson Laboratories). There was no attrition or drop out of subjects. Animal were assigned to experimental groups using simple randomization. In experiments where neutrophils were depleted, 50 µg anti-Ly6G (1A8) (BioXCell) or PBS vehicle control were delivered intraperitoneally 24 h before infection. Depletion was confirmed by flow cytometry using Ly6G (Invitrogen 367-9668-82), CD11b (Invitrogen 63-0112-82), and CXCR1 (RD Systems FAB8628P) antibodies as previously^29^. *Spy* 10^8^ CFU were suspended in 100 μL of PBS and injected subcutaneously, as previously^38^. After 24 h, mice were euthanized, lesions excised and mechanically homogenized, then stained and fixed for analysis.

### Flow Cytometry

All samples were pelleted at 12,000 x g and washed with PBS with 1 mM EDTA. For viability, samples were stained using BactoView^TM^ Dead 760/780 (Biotium cat. 40113) as per manufacturer protocol, then strained through a 100 µm filter (Avantor). Samples were fixed using 4% paraformaldehyde for 30 minutes. To stain for Group A Carbohydrate (GAC), unique to *Spy*, samples were incubated with goat anti-GAC antibody (Fitzgerald 70-XG70_R) for 1 hour, then rabbit anti-goat APC (Invitrogen A56570) for 1 hour. Samples were analyzed using BD FACSymphony A3 with excitation lasers: FITC, PE-594-A, APC, APC-Cy7. Appropriate single-color controls were used as compensation controls for these experiments. Data was analyzed using FlowJo, RRID:SCR_008520. For analysis, *speB* expressors were determined based on fluorescent intensity. Non-expressors were determined by background fluorescence from the *Spy* empty vector negative control, below 10^2^. Low expressors were identified between the ranges of 10^2^ and 10^3^, whereas high expressors were identified above 10^3^.

### Protein expression and purification

Expression plasmid pETxSUMO-Vfr was introduced to *Escherichia coli* Rosetta (DE3) and induced at 37°C for 3 hour with 1 mM isopropyl-ß-D-thiogalactopyranoside (IPTG) when O.D. 600 reached 0.4-0.6. Cells were pelleted resuspended in buffer (20 mM Tris-HCl pH 8.0) and disrupted with sonication. Lysates were centrifuged at 20,000 x g 10 minutes at 4°C. Inclusion bodies were washed twice with wash buffer 1 (20 mM Tris-HCl, 2 M urea, 0.3 M NaCl, 2% TritonX-100 pH 8.0) and dissolved in binding buffer (20 mM Tris-HCl, 0.5 M NaCl, 5 mM imidazole, 6 M guanidine hydrochloride, 1 mM 2-mercaptoethanol pH 8.0). Sample was mixed at low speed for 40 minutes at room temperature and then filter-sterilized through 0.22 µm filter (Avantor). Filtered sample was loaded to a washed and equilibrated HisTrap HP column (Cytivia) with 0.5 ml NiSO_4_. Sample within the column was washed with binding buffer and wash buffer 2 (20 mM Tris-HCl, 0.5 M NaCl, 6 M urea, 1 mM 2-mercaptoethanol pH 8.0). Refolding was performed with FPLC (AKTA start) using a urea linear gradient up to 6 M urea. Gradient volume was 30 mL and flow rate was 1 mL/minute. Sample was eluted using an imidazole linear gradient up to 400 mM imidazole in elution buffer (20 mM Tris-HCl, 0.5 M NaCl, 400 mM imidazole, 1 mM 2-mercaptoethanol pH 8.0). Gradient volume was 10 mL and flow rate was 0.5 mL/minute. Fractions containing eluted protein were pooled and quantified. SpeB was purified as previously described ^38,43^.

### Vfr Cleavage Assay

For *in vitro* cleavage experiments, recombinant Vfr (rVfr) (0.3 mg/mL) was incubated with titrations of purified SpeB (0.1 - 25 µg/mL) or recombinant Neutrophil Elastase (1 µg/mL; Sigma 324681) in assay buffer (Tris 25 mM, 300 mM NaCl, 2 mM dithiothreitol) at 37°C for 2 hours. Proteins and their cleavage products were then separated by SDS-PAGE and visualized by AcquaStain (Bulldog Bio). For neutrophil protein cleavage, purified rVfr (0.3 mg/mL) was incubated with neutrophil soluble protein (as described above) from 10^6^ cells/mL in RPMI supplemented with 2 mM dithiothreitol for 12 hours. Protease inhibitors AEBSF, Chymostatin, and PMSF were used as per manufacture protocol (G Biosciences 786-207). Neutrophil Elastase selective protease inhibitor BAY-687 (Cayman 18615) was used in titrating quantities (0.1 - 20 µg/mL).

### Protein Modeling and Cleavage Prediction

The structure of Vfr was modeled as a dimer in AlphaFold Server v3 and visualized in PyMOL, RRID:SCR_000305. Sites of potential SpeB cleavage were predicted as previously described based on known targets^44^ using ScanProsite (Expasy) with a search for the motif [IVFYM]-[ADEGKSTN] and the included condition for a net negative charge sidechain charge in the P1’-P5’ region. Sites of NET cleavage are from the motif previously described^28^.

### Protein Fractionation

Neutrophils isolated from whole blood (as above) were pelleted at 2,000 x g for 5 minutes and washed in Tris 20 mM, 1 M NaCl pH 8.0, then disrupted with sonication on ice. Neutrophil lysate was centrifuged at 20,000 x g at 4°C and filter sterilized through 0.22 µm filter (Avantor) then separated by anionic exchange by FPLC (Cytiva). Neutrophil protein was fractioned with a linear NaCl gradient up to 1 M NaCl in 20 mM of Tris. Fractions were assessed by SDS-PAGE and assayed with *Spy* containing GFP (*PspeB::gfp*) fluorescent reporter.

### Protease activity

Internally-quenched FRET peptides IFFDTWDNE, TWDNEAYVH, EAYVHDAPV, and HDAPVRSLN that detect neutrophil proteases^45^ were pooled (2.5 µM each) in PBS, 1 mM CaCl*2*, 0.01% Tween-20 (Sigma P7949) and unincubated with each column fraction. After 1 hour incubation at 37°C, the proteolysis was measured using an Nivo plate reader (PerkinElmer) with fluorophore excitation at 323 nm and emission at 398 nm as previously^45^. Measurements of SpeB activity within bacterial supernatants was performed as previously with the fluorescent peptide sub103, internally quenched with an N-terminal Mca and the C-terminal Lys-Dnp (CPC Scientific)^43^. In triplicate, 10 µM of peptide was incubated in assay buffer (PBS with 2 mM dithiothreitol) with 10 µL of supernatant at 37°C for 30 minutes. Kinetic fluorescence was measured every 30 sec (ex. 323 nm, em. 398 nm) using a Victor Nivo plate reader (PerkinElmer).

## Statistics

GraphPad Prism, RRID:SCR_002798, was used to evaluate statistical significance. Unless otherwise stated, one-way ANOVA with Dunnett’s multiple comparisons test was used for the statistical analysis of experiments and P values < 0.05 were considered significant.

## References

1. Carapetis, J. R., Steer, A. C., Mulholland, E. K. & Weber, M. The global burden of group A streptococcal diseases. Lancet Infect. Dis. 5, 685–694 (2005).

2. Misiakos, E. P. et al. Current Concepts in the Management of Necrotizing Fasciitis. Front. Surg. 1, (2014).

3. Gryllos, I. et al. Induction of group A *Streptococcus* virulence by a human antimicrobial peptide. Proc. Natl. Acad. Sci. 105, 16755–16760 (2008).

4. Tran-Winkler, H. J., Love, J. F., Gryllos, I. & Wessels, M. R. Signal Transduction through CsrRS Confers an Invasive Phenotype in Group A Streptococcus. PLoS Pathog. 7, e1002361 (2011).

5. Love, J. F., Tran-Winkler, H. J. & Wessels, M. R. Vitamin D and the Human Antimicrobial Peptide LL-37 Enhance Group A *Streptococcus* Resistance to Killing by Human Cells. mBio 3, e00394–12 (2012).

6. Brouwer, S. et al. Pathogenesis, epidemiology and control of Group A Streptococcus infection. Nat. Rev. Microbiol. 21, 431–447 (2023).

7. Carroll, R. K. & Musser, J. M. From transcription to activation: how group A streptococcus, the flesh-eating pathogen, regulates SpeB cysteine protease production. Mol. Microbiol. 81, 588–601 (2011).

8. Ikebe, T. et al. Highly Frequent Mutations in Negative Regulators of Multiple Virulence Genes in Group A Streptococcal Toxic Shock Syndrome Isolates. PLoS Pathog. 6, e1000832 (2010).

9. Ikebe, T. et al. Spontaneous mutations in Streptococcus pyogenes isolates from streptococcal toxic shock syndrome patients play roles in virulence. Sci. Rep. 6, 28761 (2016).

10. Morag R. Graham et al. Virulence control in group A *Streptococcus* by a two-component gene regulatory system: Global expression profiling and *in vivo* infection modeling. Proc. Natl. Acad. Sci. 99, 13855–13860 (2002).

11. Gryllos, I. et al. Mg^2+^ signalling defines the group A streptococcal CsrRS (CovRS) regulon. Mol. Microbiol. 65, 671–683 (2007).

12. Finn, M. B., Ramsey, K. M., Dove, S. L. & Wessels, M. R. Identification of Group A Streptococcus Genes Directly Regulated by CsrRS and Novel Intermediate Regulators. mBio 12, e01642–21 (2021).

13. Kahlenberg, J. M. & Kaplan, M. J. Little Peptide, Big Effects: The Role of LL-37 in Inflammation and Autoimmune Disease. J. Immunol. 191, 4895–4901 (2013).

14. Dorschner, R. A. et al. Cutaneous Injury Induces the Release of Cathelicidin Anti-Microbial Peptides Active Against Group A Streptococcus. J. Invest. Dermatol. 117, 91–97 (2001).

15. Schauber, J. et al. Injury enhances TLR2 function and antimicrobial peptide expression through a vitamin D–dependent mechanism. J. Clin. Invest. 117, 803–811 (2007).

16. Neely, M. N., Lyon, W. R., Runft, D. L. & Caparon, M. Role of RopB in Growth Phase Expression of the SpeB Cysteine Protease of *Streptococcus pyogenes*. J. Bacteriol. 185, 5166–5174 (2003).

17. Do, H. et al. Leaderless secreted peptide signaling molecule alters global gene expression and increases virulence of a human bacterial pathogen. Proc. Natl. Acad. Sci. 114, (2017).

18. Ma, Y., Bryant, A. E., Salmi, D. B., McIndoo, E. & Stevens, D. L. *vfr*, a Novel Locus Affecting Cysteine Protease Production in *Streptococcus pyogenes*. J. Bacteriol. 191, 3189–3194 (2009).

19. Shelburne Iii, S. A., et al. An amino-terminal signal peptide of Vfr protein negatively influences RopB-dependent SpeB expression and attenuates virulence in *Streptococcus pyogenes*. Mol. Microbiol. 82, 1481–1495 (2011).

20. LaRock, C. N. & Nizet, V. Cationic antimicrobial peptide resistance mechanisms of streptococcal pathogens. Biochim. Biophys. Acta BBA - Biomembr. 1848, 3047–3054 (2015).

21. Alberti, S., Ashbaugh, C. D. & Wessels, M. R. Structure of the *has* operon promoter and regulation of hyaluronic acid capsule expression in group A *Streptococcus*. Mol. Microbiol. 28, 343–353 (1998).

22. Blöchl, C. et al. Proteolytic Profiling of Streptococcal Pyrogenic Exotoxin B (SpeB) by Complementary HPLC-MS Approaches. Int. J. Mol. Sci. 23, 412 (2021).

23. Minns, D. et al. The neutrophil antimicrobial peptide cathelicidin promotes Th17 differentiation. Nat. Commun. 12, 1285 (2021).

24. Herwald, H. et al. M Protein, a Classical Bacterial Virulence Determinant, Forms Complexes with Fibrinogen that Induce Vascular Leakage. Cell 116, 367–379 (2004).

25. Tanaka, M. et al. Group A Streptococcus establishes pharynx infection by degrading the deoxyribonucleic acid of neutrophil extracellular traps. Sci. Rep. 10, 3251 (2020).

26. Buchanan, J. T. et al. DNase Expression Allows the Pathogen Group A Streptococcus to Escape Killing in Neutrophil Extracellular Traps. Curr. Biol. 16, 396–400 (2006).

27. Nilsson, M. et al. Activation of human polymorphonuclear neutrophils by streptolysin O from Streptococcus pyogenes leads to the release of proinflammatory mediators. Thromb. Haemost. 95, 982–990 (2006).

28. O’Donoghue, A. J. et al. Global Substrate Profiling of Proteases in Human Neutrophil Extracellular Traps Reveals Consensus Motif Predominantly Contributed by Elastase. PLoS ONE 8, e75141 (2013).

29. Doris L. LaRock, Russell, R., Johnson, A. F., Wilde, S. & LaRock, C. N. Group A Streptococcus Infection of the Nasopharynx Requires Proinflammatory Signaling through the Interleukin-1 Receptor. Infect. Immun. 88, e00356–20 (2020).

30. Hemmers, S., Teijaro, J. R., Arandjelovic, S. & Mowen, K. A. PAD4-Mediated Neutrophil Extracellular Trap Formation Is Not Required for Immunity against Influenza Infection. PLoS ONE 6, e22043 (2011).

31. Guerra, S. & LaRock, C. Group A Streptococcus interactions with the host across time and space. Curr. Opin. Microbiol. 77, 102420 (2024).

32. Kasper, K. J. et al. Bacterial Superantigens Promote Acute Nasopharyngeal Infection by Streptococcus pyogenes in a Human MHC Class II-Dependent Manner. PLoS Pathog. 10, e1004155 (2014).

33. Do, H. et al. Engineered probiotic overcomes pathogen defences using signal interference and antibiotic production to treat infection in mice. Nat. Microbiol. 9, 502–513 (2024).

34. Belaaouaj, A. Neutrophil elastase-mediated killing of bacteria: lessons from targeted mutagenesis. Microbes Infect. 4, 1259–1264 (2002).

35. Aziz, R. K. et al. Invasive M1T1 group A *Streptococcus* undergoes a phase-shift *in vivo* to prevent proteolytic degradation of multiple virulence factors by SpeB. Mol. Microbiol. 51, 123–134 (2004).

36. Kansal, R. G., Nizet, V., Jeng, A., Chuang, W.-J. & Kotb, M. Selective Modulation of Superantigen-Ind Responses by Streptococcal Cysteine Prot.

37. Lauth, X. et al. M1 Protein Allows Group A Streptococcal Survival in Phagocyte Extracellular Traps through Cathelicidin Inhibition. J. Innate Immun. 1, 202–214 (2009).

38. LaRock, D. L. et al. Group A Streptococcus induces GSDMA-dependent pyroptosis in keratinocytes. Nature 605, 527–531 (2022).

39. Bjånes, E. et al. An efficient *in vivo*-inducible CRISPR interference system for group A *Streptococcus* genetic analysis and pathogenesis studies. mBio 15, e00840–24 (2024).

40. Klock, H. E. & Lesley, S. A. The Polymerase Incomplete Primer Extension (PIPE) Method Applied to High-Throughput Cloning and Site-Directed Mutagenesis. in High Throughput Protein Expression and Purification (ed. Doyle, S. A.) vol. 498 91–103 (Humana Press, Totowa, NJ, 2009).

41. Jeng, A. et al. Molecular Genetic Analysis of a Group A *Streptococcus* Operon Encoding Serum Opacity Factor and a Novel Fibronectin-Binding Protein, SfbX. J. Bacteriol. 185, 1208–1217 (2003).

42. Wilde, S. et al. Detoxification of reactive oxygen species by the hyaluronic acid capsule of group A *Streptococcus*. Infect. Immun. 91, e00258–23 (2023).

43. LaRock, C. N. et al. IL-1β is an innate immune sensor of microbial proteolysis. Sci. Immunol. 1, (2016).

44. Johnson, A. F. et al. Proinflammatory synergy between protease and superantigen streptococcal pyogenic exotoxins. Infect. Immun. 93, e00405–24 (2025).

45. Sun, J. et al. The Pseudomonas aeruginosa protease LasB directly activates IL-1β. EBioMedicine 60, 102984 (2020).

